# Transient ATR inhibition following ionizing radiation enhances immune-mediated antitumor response and survival

**DOI:** 10.64898/2026.05.25.727700

**Authors:** Joshua Deppas, Brian F. Kiesel, Frank P. Vendetti, Pinakin Pandya, Jianxia Guo, Kristine L. Cooper, Matthew J. Bakkenist, Meysam Tavakoli, Maria diMayorca, Naveed M Islam, David A. Clump, Christopher J. Bakkenist, Jan H. Beumer

**Affiliations:** Cancer Therapeutics Program, UPMC Hillman Cancer Center, University of Pittsburgh, Pittsburgh, Pennsylvania, USA; Department of Pharmaceutical Sciences, School of Pharmacy, University of Pittsburgh, Pittsburgh, Pennsylvania, USA; Department of Radiation Oncology, UPMC Hillman Cancer Center, School of Medicine, University of Pittsburgh, Pittsburgh, Pennsylvania, USA; Biostatistics Facility, UPMC Hillman Cancer Center, University of Pittsburgh, Pittsburgh, Pennsylvania, USA; Department of Radiation Oncology, Winship Cancer Institute, Emory University, Atlanta GA, USA; Department of Radiation Oncology, School of Medicine, West Virginia University, Morgantown, West Virginia, USA; Department of Pharmacology and Chemical Biology, UPMC Hillman Cancer Center, School of Medicine, University of Pittsburgh, Pittsburgh, Pennsylvania, USA; Department of Oncology, Johns Hopkins University School of Medicine and Johns Hopkins Sidney Kimmel Cancer Center, Baltimore, Maryland, USA

**Keywords:** ATR, ceralasertib, elimusertib, berzosertib, radiation

## Abstract

**Background:** ATR activation following DNA damage from cancer treatments such as radiation can mitigate anticancer efficacy, making ATR inhibitors (ATRi) an attractive therapeutic. *In vivo* and *in vitro* studies have shown enhanced tumor cell radiosensitivity with the ATRi ceralasertib, elimusertib, and berzosertib, however, the potentiating effect of ATRi on ionizing radiation (IR) through immune-based mechanisms has only been studied with ceralasertib.

**Methods:** We aimed to determine if antitumor immune responses observed with ceralasertib in combination with IR extend to the other ATRi class members in the preclinical CT26 mouse model. We also examined the relationship between exposure and immune stimulation, efficacy and survival outcomes of each ATRi when combined with IR.

**Results:** Ceralasertib and elimusertib, not berzosertib, synergized with IR in a dose and schedule-dependent manner to modify tumor antigen-specific CD8^+^ T cell populations in the draining lymph node. Transient ATRi therapy, combined with IR, enhances antitumor efficacy, promoted tumor shrinkage, and increased survival. ATRi elicited differential inflammatory gene induction and dose-dependent unique cytotoxicity profiles *in vitro*.

**Conclusion:** The immune mediated antitumor effect of ATRi combined with radiation is dose and schedule dependent, and while likely a class effect, may differ between ATRi compounds.

## 1 INTRODUCTION

Cancer cells survive DNA damage by activation of the DNA damage response (DDR), a highly organized and coordinated network of signaling cascades that promote DNA repair through cell-cycle checkpoint activation. Apical to the DDR is the ataxia telangiectasia and Rad3-related (ATR) protein kinase. ATR activation induced by DNA damaging cancer treatments can mitigate their anticancer effects, thus making ATR an attractive therapeutic target for inhibition.

Several ATR inhibitors (ATRi) have been developed, including: berzosertib (VX-970, M6620), ceralasertib (AZD6738), elimusertib (BAY-1895344), and more recently tuvusertib (M1774) and camonsertib (RP-3500). These agents potentiate the effects of ionizing radiation (IR) and other DNA damaging agents and kill cell lines with mutations in DDR proteins (1–5). Furthermore, emerging literature shows that ATR shapes the immune landscape in the tumor microenvironment and periphery, as ATRi increase tumor MHC-I expression, reduce tumor PD-L1 expression, potentiate IR-induced IFN-I signaling, and generate durable immunological memory in preclinical models (6–10).

Although *in vitro* studies have shown enhanced tumor cell radiosensitivity with ceralasertib (11), elimusertib (12), berzosertib (13, 14) and its predecessor, VE-821 (15–17), only ceralasertib has been studied in *in vivo* studies examining the potentiating effect of ATRi on IR through immune-based mechanisms (8, 10, 11, 18, 19). In mice ceralasertib, in combination with IR, has been shown to increase tumor-infiltrating cytotoxic T lymphocytes (CTL) as well as tumor antigen-specific CTLs in the tumor-draining lymph node (DLN) (10, 19). Furthermore, only a single dose level of ceralasertib (75 mg/kg) has been evaluated in this manner, and it is therefore unknown what minimum exposure of ATRi may be needed to elicit these immunostimulatory effects. As additional ATRi progress through development it is important to identify if these immune effects observed with ceralasertib extend to the entire drug class.

Here, we show that synergistic murine antitumor immune responses, originally observed with ceralasertib in combination with IR, extend to elimusertib, but not berzosertib. We further show that stepwise increases to clinically-relevant ATRi dose modulate tumor antigen-specific CD8^+^ T cells in the DLN in a dose-dependent manner. Finally, we show that, when combined with IR, transient ATR inhibition is required for tumor antigen-specific CD8^+^ T cell expansion in the DLN and that the immunostimulatory enhancement associated with ceralasertib and elimusertib translate to increased survival in CT26 syngeneic tumor models.

## 2 RESULTS

### ATRi synergizes with IR in a dose-dependent manner to modify tumor antigen-specific CD8^+^ T cell populations in the DLN

Short-course ceralasertib administration on days 1-3 (QDx3) synergizes with 2 Gy IR on days 1-2 (2 Gy IR x 2) to increase the population of functional tumor-infiltrating CD8+ T cells (i.e., CTLs) by days 9-12 (10). Later studies show that the same ceralasertib regimen integrates with IR to generate an expansion of peripheral tumor antigen-specific CD8+ T cells in the DLN by day 9 (19). Evidence for this phenomenon is currently limited to a single member of the ATRi class administered at only a single dose (ceralasertib at 75 mg/kg) and the optimal ATRi dose needed for maximal response is unclear. Previous PK studies of ATRi revealed widespread, non-linear distribution to various tissues, complicating predictions of antitumor activity in relevant sites such as DLN (20–22). We hypothesized that, when combined with IR, the expansion of tumor-specific CD8+ T cells is ATRi dose-dependent and similar class-effects will be observed with alternative ATRi.

A range of doses of ceralasertib, elimusertib, or berzosertib (administered on days 1-3) combined with 2 Gy IR (administered on days 1-2), were applied to compare the response of tumor antigen-specific CD8+ T cells in the DLN and tumor on day 9 (**Figure 1A**). Tumor-bearing female BALB/c mice were stratified to receive IR alone, ATRi (ceralasertib, elimusertib, or berzosertib) alone, IR in combination with one of four increasing ATRi doses, or respective vehicle controls. The doses for ceralasertib were: 2, 7.5, 20, or 75 mg/kg. The doses for elimusertib were: 1, 4, 10, or 40 mg/kg. The doses for berzosertib were: 2, 6, 20, or 60 mg/kg. On day 9, the tumor and DLN were removed and CD8^+^ T cells were labeled with an MHC class I H-2Ld-restricted Pentamer loaded with AH1 peptide, the immunodominant CT26 antigen, and detected using flow cytometry (**Figure 1B)**.

**Figure 1.**
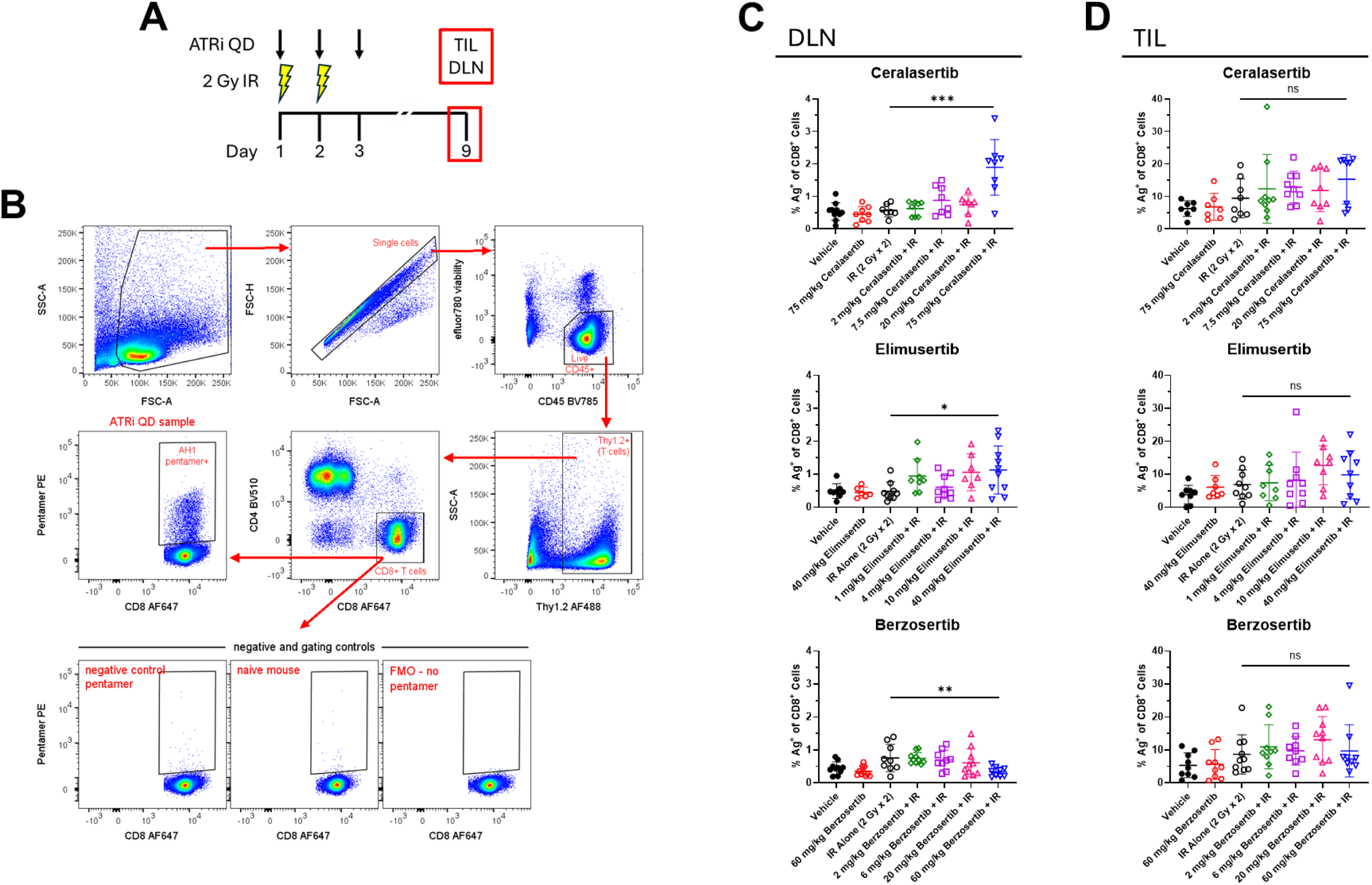
The A) treatment schema that was followed for the radiation dose range finding studies and B) flow cytometry gating strategy with representative cytograms depicting Pentamer+ CD8+ T cells. Fluorescence-minus-one (no Pentamer), naïve (negative, no tumor), and negative Pentamer controls shown. Composition of CD8^+^ T cells that are CT26 antigen specific for all groups in the C) draining lymph node (DLN) and D) tumor (TIL). Data represented as mean (bold line) and standard deviation. *\*P < 0.05, **P < 0.01, ***P < 0.001*, by fitting data to a linear model across the treatment dosing range (see Suppl.Fig. 1). NS: not significant.

In the DLN, a positive association was observed between ATRi dose and tumor antigen-specific CTL levels, as determined by a linear model for ceralasertib (*p<0.001*) and elimusertib (*p=0.030*) combination cohorts (**Figure 1C and Suppl.Fig. 1**), indicating that antigen-specific CD8^+^ T cell counts increased in a dose-dependent manner. In contrast, a statistically significant negative association was observed between berzosertib dose and antigen-specific CD8^+^ T cell levels in IR combination cohorts (*p=0.001*), suggesting a dose-dependent immunosuppressive effect with berzosertib (**Figure 1C**). We did not observe a significant change in tumor antigen-specific CD8^+^ T cells in the tumor infiltrating lymphocytes (TILs) at this timepoint for any ATRi combination treatment cohorts (**Figure 1D and Suppl.Fig. 1**). Although changes to antigen-specific CD8^+^ T cells were observed in the DLN with each ATRi cohort, these observations were not associated with any non-specific changes to CD45^+^CD8^+^ immune cells (**Suppl.Fig. 2A-C**), suggesting an overall stable cytotoxic T cell population. Ultimately, while elimusertib mirrored ceralasertib in having a dose-dependent immunostimulatory effect, berzosertib did not, suggesting this is not a class effect extending to all ATRi.

### Dosing ceralasertib twice a day in combination with IR abolishes tumor antigen-specific CTL expansion in the DLN

We have previously shown that berzosertib has a half-life approximately 3-fold longer than ceralasertib or elimusertib in CT26 tumor-bearing female BALB/c mice (6.12 h vs 1.75 and 1.80 h, respectively) (20–22). While short-term ATR inhibition is known to increase tumor immunogenicity, upregulate type I IFNs and chemokines, enhance immune cell infiltration, and sensitize tumors to radiation therapy, persistent inhibition has been shown to suppress T cell proliferation, impair hematopoietic stem cell function, and exhaust immune cells (1, 6, 10, 19, 23, 24). Specifically, recent studies have shown that 9 days of once-daily ceralasertib treatment abolishes peripheral expansion of tumor antigen-specific CD8^+^ T cell responses following IR. We hypothesized that such detrimental effects were also induced by berzosertib owing to its long half-life. To phenocopy berzosertib’s long half-life, we conducted experiments to compare once-daily (QD) *versus* twice-daily (BID) dosing of ceralasertib or elimusertib in combination with IR. CT26 tumor-bearing mice were stratified to receive vehicle controls, IR alone (2 Gy x 2), 75 mg/kg ceralasertib or 40 mg/kg elimusertib alone or in combination either QD, resembling the regimen used previously, or BID on days 1-3, with IR on days 1-2. As our previous data provided no evidence to suggest any significant change to AH1 antigen-specific CTLs in TILs on day 9, only the DLN was collected.

Initially, BID dosing of either ceralasertib or elimusertib yielded lower, but statistically non-significant, antigen-specific CD8^+^ T cell counts than after QD dosing (**Suppl.Fig. 3A,B and Suppl.Table 1**). Between-study comparison of the DLN suggested suppressed antigen-specific CD8^+^ T cells in both IR alone cohorts (average: 0.314%) compared to our previously generated data (average: 0.523%). We believed our positive control, QD ATRi in combination with IR, failed to provide a biologically accurate result against which to appropriately compare the BID ATRi in combination with IR cohorts, as all treatment cohorts appeared to be suppressed, similar to the IR alone group. We attributed this loss of signal to the manner in which mice were irradiated. Mice received 2 Gy radiation (6mV photon energy, 2 cm field) while immobilized on a clinical radiation treatment table, with tumors facing at an upward angle. While this technique has demonstrated effectiveness (19), variability in precision while targeting the tumor is inherent, and irradiating the DLN is also likely.

Following these experiments, and with the installation and availability of an image-guided small animal irradiator, we modified our treatment protocol such that mice would be anesthetized with isoflurane and receive 2 Gy conformal radiation via a 10 mm collimator and Cu treatment filter using a Precision SmART+ (225 kV) small animal irradiator. This method has proven to be safe with no anticipated drug-drug interactions with isoflurane and ATRi (19). Since ceralasertib in combination with IR provided the most pronounced increase of antigen-specific CD8^+^ T cells, we chose to repeat our ATRi QD vs BID in combination with IR experiments with this ATRi and incorporate the new, targeted IR treatment protocol. CT26 tumor-bearing mice were stratified to receive IR alone (2 Gy x 2), QD 75 mg/kg ceralasertib + IR, or BID 75 mg/kg ceralasertib + IR in a similar manner as previously described (**Figure 2A**) and DLNs were examined for tumor antigen-specific CD8+ T cells on day 9 (**Figure 2B)**.

**Figure 2.**
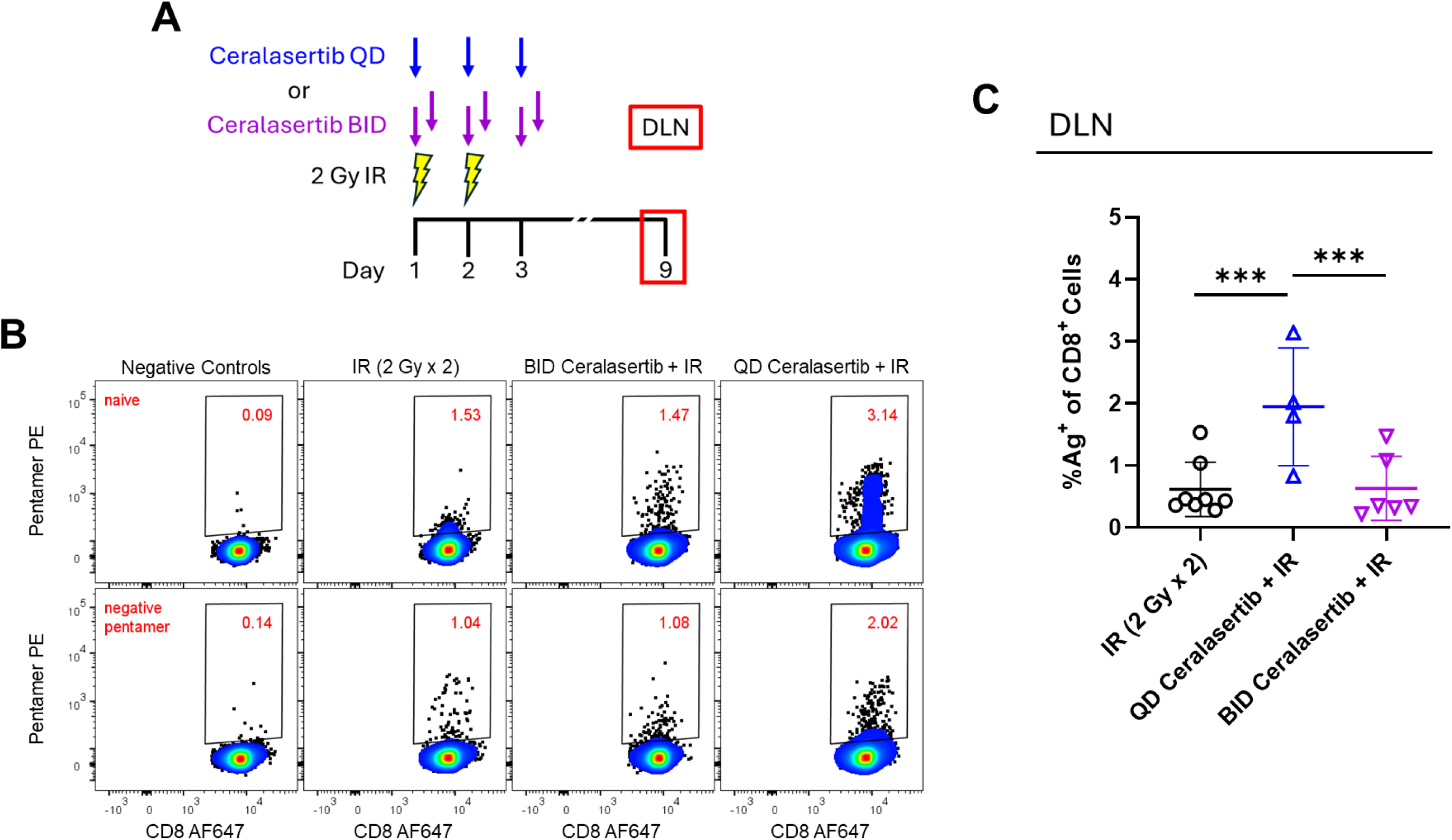
The A) treatment schema that was followed for the ceralasertib QD vs BID studies and B) representative cytograms depicting Pentamer^+^ CD8^+^ T cells, naïve (negative, no tumor), and negative pentamer controls shown. C) Composition of CD8^+^ T cells that are CT26 antigen specific in the draining lymph node. Data represented as mean (bold lines) and standard deviation. *\*\*\*P < 0.001* by a generalized linear model. Brackets not shown were not statistically significant.

Analysis of the DLN showed that QD 75 mg/kg ceralasertib in combination with IR significantly increased the proportion of tumor antigen-specific CD8^+^ T cells compared to the BID ceralasertib combination cohort (1.95% vs 0.630% antigen-specific CD8+ T cells; *p<0.001*) and the IR alone controls (1.95% vs 0.615% antigen-specific CD8+ T cells; *p<0.001*) (**Figure 2C and Suppl.Table 2**). In contrast, BID ceralasertib administration in combination with IR abrogated tumor antigen-specific T cell expansion, resulting in similar average compositions to that of mice receiving IR alone. Collectively, these results suggest that brief, transient ATR inhibition, when combined with short course IR, achieves antigen-specific CTL expansion in the DLN.

### Transient ATRi administration combined with IR slows tumor growth, increases survival, and is associated with anti-tumor memory

To assess the efficacy of the addition of ceralasertib, elimusertib, or berzosertib to IR, we retained the QDx3 ATRi treatment regimen used to elicit immune responses with ceralasertib and elimusertib and repeated it after two weeks. The time between ATRi cycles was considered long enough to allow for proper immune rebound while minimizing unchecked tumor growth (25). CT26 tumor-bearing mice were stratified, primarily by tumor size and secondarily by total body weight (**Suppl.Fig. 4A,B**), to receive either IR alone or in combination with ceralasertib (75 mg/kg), elimusertib (10 mg/kg), or berzosertib (20 mg/kg). This regimen incorporated two cycles of the previously described ATRi (QDx3) and IR (2 Gy x 2) treatment regimen and allowed ATRi and IR to be administered over the course of 4 weeks, with 3 days on/11-day-off serving as a single cycle (i.e., ATRi on days 1-3 and 15-17 and IR on days 1-2 and 15-16) (**Figure 3A**). Mice receiving IR were exposed to 2 Gy tumor targeted radiation using an image-guided small animal irradiator as previously described.

**Figure 3.**
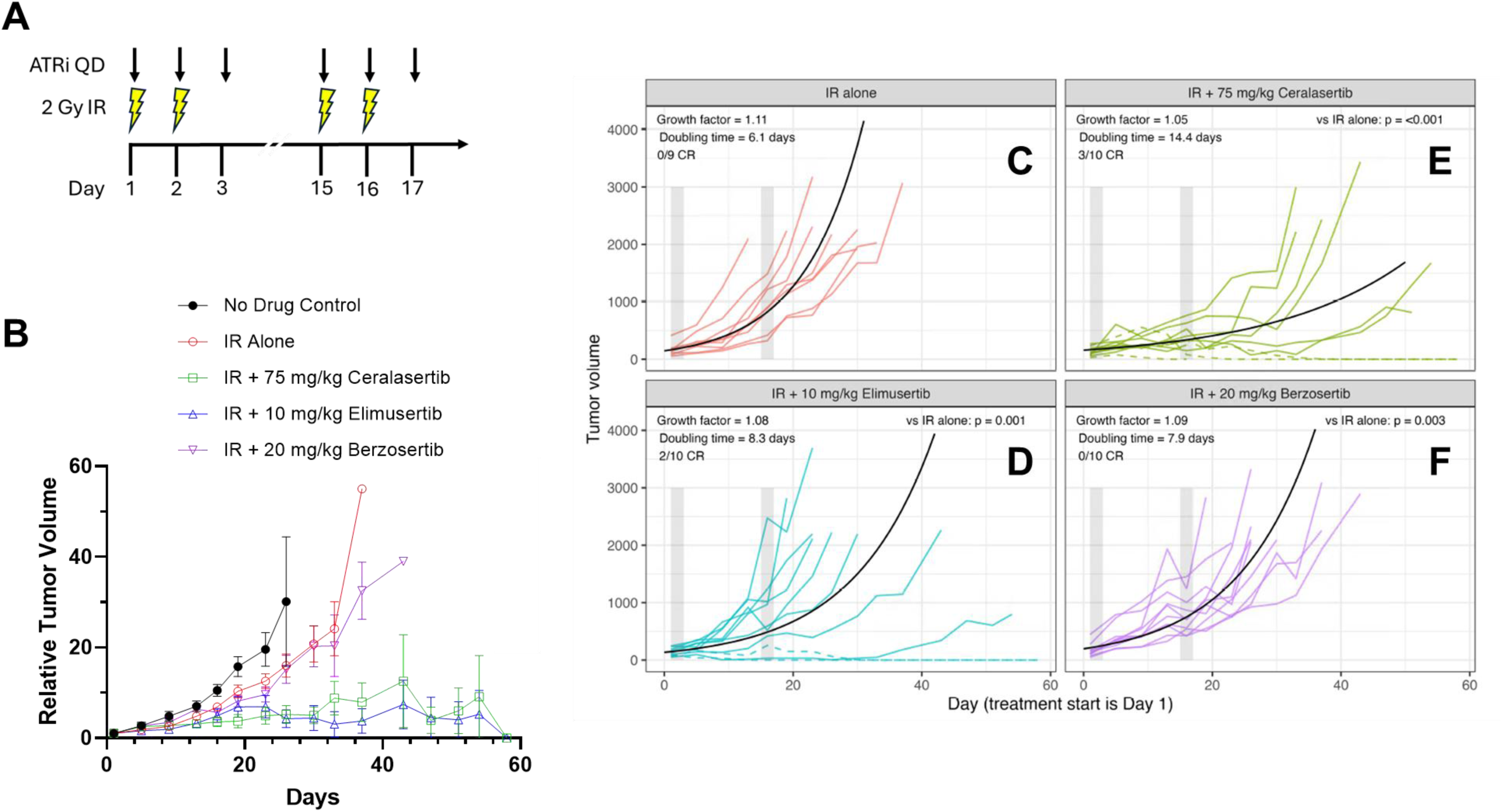
A) treatment schema that was followed for IR +/- ATRi efficacy and survival study. B) relative tumor volume from day 1. C) tumor growth dynamics depicting a linear mixed-model fit to adjust for repeated measures over time. Data represented as mean (bold line) and SEM. Black lines represent estimated growth curves for each group (excluding complete responders). Mice that are complete responders are represented by “dashed” lines. Pairwise comparisons were made to the IR alone control group. Shaded vertical gray bars indicate treatment days with ATRi ± IR.

Day 26 was chosen as an acceptable experimental timepoint for comparisons with combination treatment cohorts, as it represented a time approaching the end of the two-cycle regimen. At this time, half of the mice treated with IR alone were euthanized for exceeding acceptable tumor size, while the other half harbored tolerable tumors. This compared to only two surviving mice with tolerable tumors in the no drug control cohort, suggesting modest antitumor activity from IR alone. Monitoring of individual tumor growth showed that the addition of berzosertib to IR had limited impact on tumor progression compared to controls (**Suppl.Fig. 5A**). Conversely, the addition of ceralasertib or elimusertib to IR resulted in observable tumor suppression and delay, with one mouse from each cohort achieving complete regression (**Suppl.Fig. 5A**). Between-group analysis of the percent change in tumor volume on day 26 relative to day 1 confirmed a statistically significant reduction in tumor burden in mice receiving either ceralasertib (*p=0.023)* or elimusertib (*p=0.020)* in combination with IR (**Suppl.Fig. 5B**). Combination treatment with berzosertib resulted in no visible change from IR alone (*p=0.992*). These findings further support the thesis that transient ATR inhibition achieved with ceralasertib or elimusertib, but not the continuous ATR inhibition with berzosertib, enhances immunologic antitumor activity after IR.

To assess the long-term therapeutic benefit of ATRi combined with IR, we extended our analysis to further evaluate tumor shrinkage, survival, and exploratorily evaluate potential immunological consequences of our transient ATRi treatment model. By day 58, mice in all treatment cohorts were either euthanized for exceeding acceptable tumor size or had responded to treatment resulting in complete tumor clearance. Mice in the IR alone cohort had consistent and rapid tumor progression, and ultimately experienced a 55-fold increase from baseline tumor volume with a tumor doubling time of 6.1 days (**Figure 3B,C**). Following analysis of growth curves, ceralasertib (*p<0.001*), elimusertib (*p=0.001*), and berzosertib (*p=0.003*) combination cohorts were all found to significantly improve tumor shrinkage over time compared to IR treatment alone (**Figure 3C-F**). The elimusertib cohort comprised of two complete responders (mice achieving complete tumor clearance) with an associated tumor doubling time of 8.3 days (**Figure 3D)**. The ceralasertib cohort comprised of three complete responders with an associated tumor doubling time of 14.4 days (**Figure 3E**). However, in one mouse a complete response was documented for one week followed by dosing-related complications and death during the second treatment cycle. One mouse in each of the elimusertib and ceralasertib cohorts displayed tumor growth of 4.1 and 2.6 times baseline tumor volumes, respectively, prior to achieving a complete tumor clearance. Mice receiving berzosertib in combination with IR demonstrated a mean relative tumor increase of 39 and a tumor doubling time of 7.9 days (**Figure 3B,F**). Individual mouse tumor volumes for each cohort can be seen in (**Suppl.Fig. 6A-E**).

The observed antitumor effects aligned with long-term survival analyses (**Figure 4A and Suppl.Table 3**). Mice in the no drug control and IR alone groups exhibited maximum survival endpoints at days 26 and 37, respectively. The addition of berzosertib to IR resulted in a slightly longer maximum survival endpoint of 43 days. Overall survival was longer in mice receiving IR plus ceralasertib or elimusertib, as two mice in each of these combination cohorts achieved complete tumor response by completion of the study (day 58). To test statistical differences in overall survival, a Cox proportional hazards model was fit for all combination cohorts. Analysis found that IR plus ceralasertib or elimusertib resulted in significantly longer survival outcomes compared to IR alone (**Suppl.Table 4**). As these were the only two cohorts to contain apparent complete responders, we next investigated the role of CD8^+^ T cells in maintaining tumor control. After a tumor-free period of at least 10 days, CTLs from apparent complete responders were depleted by dosing 250 µg anti-CD8 antibody (αCD8) IP QDx3 (days 48, 51, and 54) and monitored for tumor regrowth (**Figure 4B**). Neither cohort resulted in tumor regrowth, however mice continued to be monitored for any changes to behavior and overall health.

**Figure 4.**
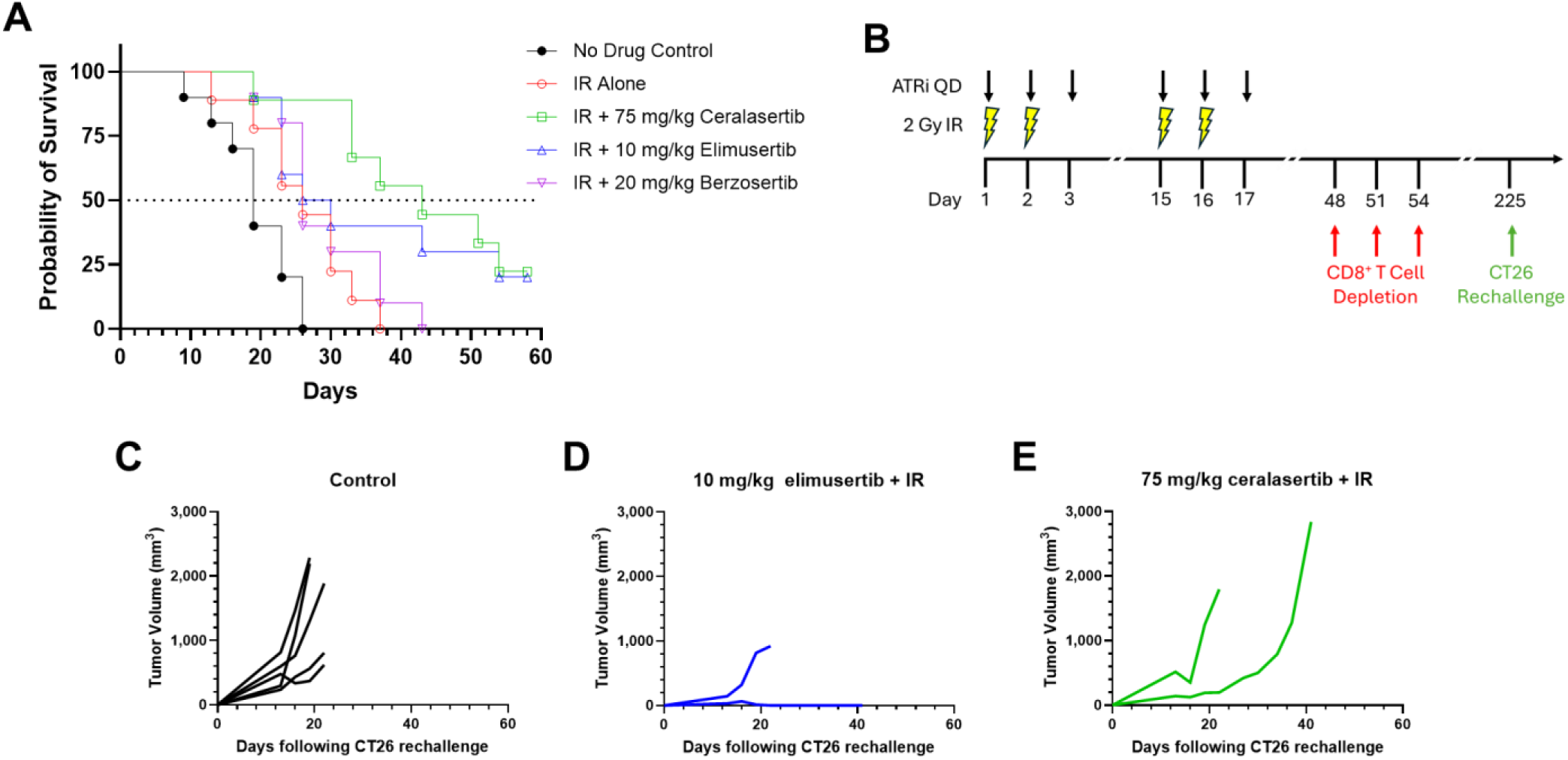
The A) probability of survival in all treatment groups during the entirety of the IR +/- ATRi efficacy and survival study. The B) schema that was followed to CD8-deplete mice on days 48, 51, and 54 and, following no observed tumor re-growth, rechallenge with CT26 cells and day 225 C-E) absolute tumor volume in complete responders and control mice following tumor rechallenge.

Following a tumor-free period of approximately 170 days, the complete responder mice that had previously undergone CD8+ T cell depletion, were rechallenged with CT26 cells, parallel to previously tumor-naïve BALB/c mice. Tumor-naïve BALB/c control mice (N=5) each developed tumors (**Figure 4C**). Each complete responder was able to establish CT26 tumors upon rechallenge, however most tumor re-growth appeared to be slower compared to the control mice. In the elimusertib cohort, one mouse achieved complete loss by day 27 post-rechallenge after initial tumor take and growth to a maximum of 63 mm^3^ (**Figure 4D**). The other mouse survived for 22 days post rechallenge. In the ceralasertib cohort, one mouse experienced noticeably delayed tumor growth resulting in survival up to day 41 post-rechallenge (**Figure 4E**). The second mouse displayed tumor regrowth similar to that of the parallel control mice.

### ATRi elicit differential inflammatory gene induction and cytotoxicity in vitro

It has previously been shown that ATRi treatment integrates with IR to affect the chemokine environment in the tumor microenvironment (TME) post IR. The dependence on IFN-β of immune responses after IR is well established, and increases of IFN-β, CXCL10, and CCL2 within tumors are known to affect immune cell recruitment and activation (26, 27). IFN-β signaling has been shown to regulate natural killer (NK) cell function and T cell activation while also inducing the production of: CXCL10, which recruits antitumor NK cells and effector T cells; and CCL2, ultimately triggering the recruitment of inflammatory monocytes (26–29). To compare immunogenic and cytotoxic effects of ATRi, CT26 cells were treated *in vitro* for 24 h using the previously defined IC^25^ concentrations of ceralasertib, elimusertib, and berzosertib, respectively (25). Gene expression was measured in untreated and ATRi-treated cells via qPCR analysis for the pro-inflammatory cytokine, IFN-β, and pro-inflammatory chemokines, CXCL10 and CCL2. Results were obtained from two independent, technical replicate experiments. Numerically, IFN-β was upregulated following treatment with all 3 ATRi, however only elimusertib generated a statistically significant induction (*p=0.002*) (**Figure 5A**). CCL2 gene expression values increased non-significantly with elimusertib (*p = 0.133*). CCL2 gene expression with ceralasertib was similar to controls, while after berzosertib CCL2 gene expression values decreased non-significantly (*p=0.539*) (**Figure 5B**). All three ATRi demonstrated higher CXCL10 expression values compared to controls, however only with elimusertib did this reach statistical significance (*p<0.001*) (**Figure 5C**). These findings suggest a selective activation of specific innate immune signaling pathways in response to ATRi-induced DNA damage. The accumulation of DNA lesions following ATRi treatment results in cytosolic DNA fragments, which in turn activates the cGAS-STING pathway leading to the phosphorylation of TBK1 and IRF3 and subsequent transcriptional activation of type I IFNs, including IFN-β (30). IFN-β then stimulates JAK-STAT signaling to increase expression of interferon-stimulated genes (ISGs), such as CXCL10. In contrast, CCL2 is uniquely regulated by the transcription factors NF-κB and AP-1, which are typically activated by pro-inflammatory stimuli such as TNF-α or oxidative stress. As previous reports have shown CCL2 induction following ATRi in combination with IR, the lack of CCL2 expression change here suggests that ATR inhibition alone may not be sufficient enough to engage these pathways (19).

**Figure 5.**
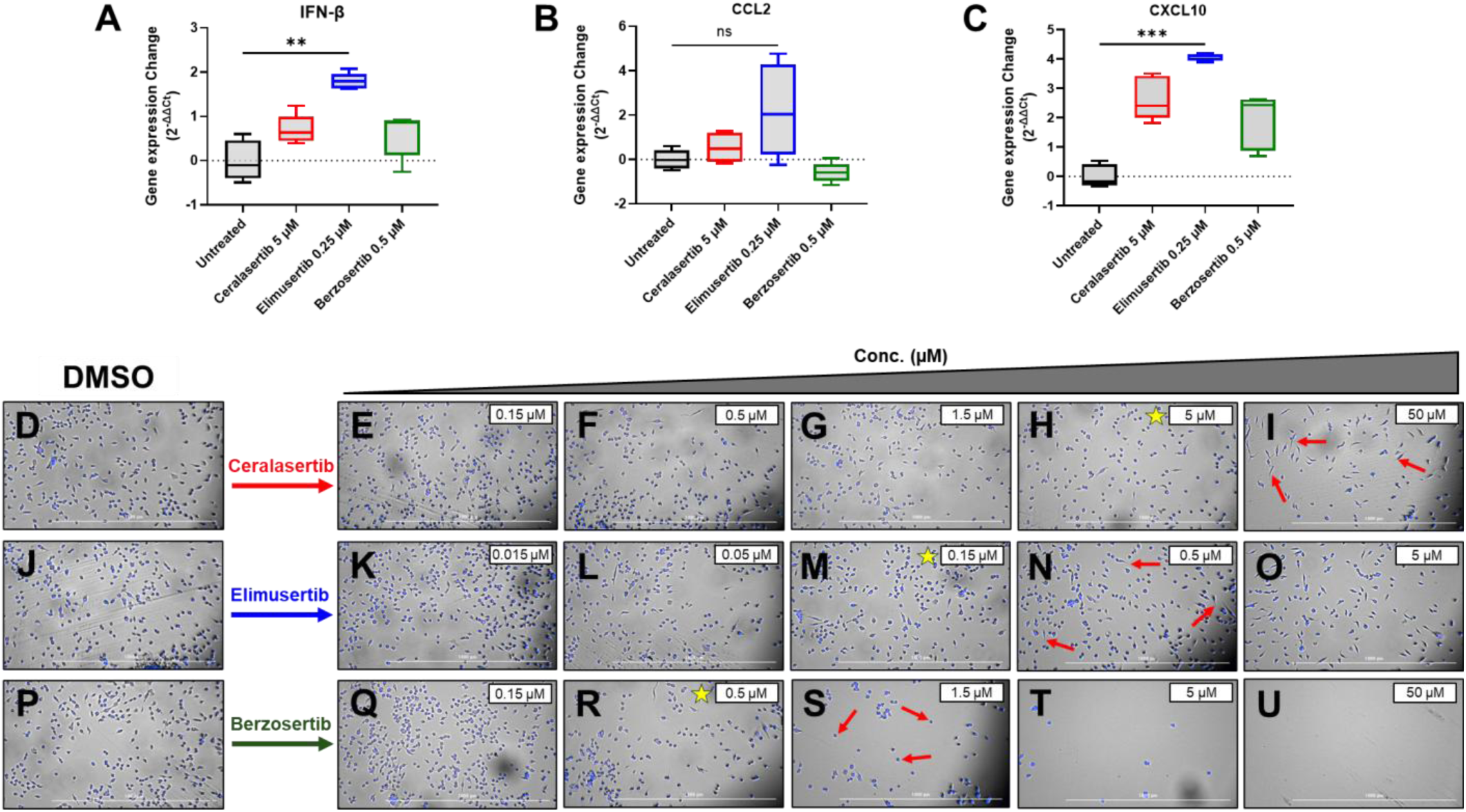
Gene expression data representing two separate experiments of A) IFN-β B) CCL2 and C) CXCL10 in CT26 cells at 24 h following exposure to IC^25^ concentration levels of ceralasertib, elimusertib, and berzosertib. Representative cellular morphology of CT26 cells 24 h after exposure to DMSO (D, J, and P) and ceralasertib E) 0.15 µM, F) 0.5 µM, G) 1.5 µM, H) 5 µM, and I) 50 µM. Elimusertib K) 0.015 µM, L) 0.05 µM, M) 0.15 µM, N) 0.5 µM, and O) 5 µM. Berzosertib Q) 0.15 µM, R) 0.5 µM, S) 1.5 µM, T) 5 µM, and U) 50 µM. Yellow stars denote IC^25^ (or closest) concentration. Red arrows represent distinct cellular morphology. *\*\*P < 0.01, ***P < 0.001*, by Friedman test with Dunn’s multiple comparisons test with untreated cells serving as the control.

To evaluate morphological changes, CT26 cells were treated with increasing concentrations of ceralasertib (0.15, 0.5, 1.5, 5, and 50 µM), elimusertib (0.015, 0.05, 0.15, 0.5, and 5 µM), or berzosertib (0.15, 0.5, 1.5, 5, and 50 µM) for 24 h. Brightfield microscopy was used to visualize any ATRi-induced morphological changes. Compared to DMSO-treated controls, cells exposed to ATRi exhibited concentration-dependent cell death and alteration to their morphology. Changes to cellular morphology included increased elongation and tail-like extensions with respect to concentration for ceralasertib (**Figure 5D-I**) and elimusertib (**Figure 5J-O**). On the other hand, berzosertib treated cells displayed more unique characteristics that consisted of a compact, contracted cell shape and rounded morphology (**Figure 5P-U**).

These results suggest ATRi elicit distinct and unique immunomodulatory and cytotoxic effects. The compound-specific morphological changes further suggest divergent mechanisms of action between ceralasertib/elimusertib *versus* berzosertib, which may influence both direct tumor cell killing and the impact of immune cell recruitment to the TME.

## 3 DISCUSSION

Our findings support the development of targeted therapeutics capable of synergizing with IR to directly enhance tumor cell death and simultaneously activate immune responses to increase antitumor benefit. In studies presented here, we demonstrate that multiple ATRi can synergize with IR in a dose-dependent manner to modify tumor antigen-specific CD8+ T cell populations. Our results confirm previous observations for ceralasertib (19) and expand this finding to show elimusertib also synergizes with IR to increase tumor immunogenicity. When combined with IR, QD dosing of shorter half-life ATRi, ceralasertib and elimusertib, promoted CD8^+^ T cell expansion in the DLN. In contrast, the longer half-life ATRi, berzosertib, suppressed CD8^+^ T cells in the DLN. Similar CD8^+^ T cell suppression was also observed with BID ceralasertib, suggesting that ATR inhibition is indeed required to be transient for immune stimulation, and thus not achievable with berzosertib. When combined with IR, all 3 ATRi significantly reduced tumor burden, but only the addition of ceralasertib and elimusertib significantly prolonged survival, further separating berzosertib from its other class members. A similar discrepancy pattern between ATRi was also observed with cellular morphology, suggesting an inconsistent mechanistic class effect and further supporting immune-mediated tumor cell death associated with ceralasertib and elimusertib and direct tumor cell killing with berzosertib.

Previous work has shown that proliferating T cells are transiently depleted by short-course treatment (QDx3) with ATRi and cessation of this therapy is required for rebound and expansion by day 7 (i.e., 4 days after cessation) (10, 19). However, these findings were limited to only a single 75 mg/kg dose level of ceralasertib. Here, using a similar dosing regimen, we show dose-dependent expansion of CD8^+^ T cells following treatment with both ceralasertib and elimusertib, with the highest tested doses being 75 mg/kg and 40 mg/kg, respectively. At these doses, we have previously determined plasma exposure (i.e., AUC) to be more than 7-fold higher for ceralasertib compared to elimusertib (20, 21), yet, in our present studies, immunostimulatory effects were observed with both agents. Therefore, target site exposure may better explain the observed immune responses and, although tumor partitioning was similar, elimusertib exhibited approximately 2-fold higher distribution into the DLN. Thus, it is plausible that higher DLN exposure, combined with higher overall potency (25), acutely induced greater T cell depletion during treatment, creating the niche into which the expansion of tumor antigen-specific CD8^+^ T cells can occur, and may ultimately be a contributing factor to why elimusertib experienced similar responses to ceralasertib despite lower systemic exposure (31–33).

Emerging evidence suggests that prolonged inhibition or disruption of DDR pathways may also impair adaptive immune functions, particularly within the TME and DLN (31). Although aberrations in the DDR are not known to directly affect antigen recognition, downstream consequences can be highly dependent on DDR status. Particularly during T cell activation and clonal expansion, where rapid proliferation imposes substantial replication stress that necessitates intact DDR signaling for genomic stability and cell survival (32–34). Therefore, the duration of ATR inhibition is likely critical for optimizing combination with DNA damaging therapies as sufficient time following ATR inhibition with ceralasertib is needed to generate a niche environment within the tumor and DLN into which new, tumor antigen-specific CTLs can expand (19). Furthermore, the administration of ceralasertib daily for 9 consecutive days (QDx9) has been shown to abolish the expansion of CTLs in the DLN and impair clonal expansion of activated T cells. Here, berzosertib (QDx3) administration mirrored this abolishment and was uniquely associated with dose-dependent suppression of tumor antigen-specific CD8^+^ T cells. Previous PK studies have determined elimination half-lives of 1.8, 1.8, and 6.1 h, for ceralasertib, elimusertib, and berzosertib, respectively, indicating a more than 3-fold persistence of ATR inhibition for berzosertib (20–22). We hypothesized that the absence of antigen-specific CD8^+^ T cell expansion observed with berzosertib could be due to its longer half-life. By administering ceralasertib BID, we were able to recapitulate the effects of berzosertib, and phenocopy the absence of tumor-specific CD8^+^ T cell expansion. Measurement of tumor antigen-specific CD8^+^ T cells in the TILs yielded no significant changes between IR alone and combination with any ATRi, suggesting a temporal or spatial disconnect in the immune response observed in the DLN. However, since our data clearly shows antitumor activity with this schedule, the most likely scenario is that we missed the CD8^+^ T cell signal in the tumor due to the time of our sampling on day 9. Prior studies demonstrated continued infiltration of CT26 tumors by functional CTLs between day 9 and day 12 (10), supporting that our day 9 sampling was too early to realize differences in AH1 antigen-specific CTLs in the tumors.

In line with our immunological findings, our efficacy analysis found differences between ATRi when combined with IR. The significant expansion of CD8^+^ T cells observed with ceralasertib and elimusertib translated to enhanced antitumor activity and extended survival. Although berzosertib was associated with immune suppression, efficacy studies showed significant tumor growth inhibition and non-significant increases in overall survival. Unique to berzosertib, our *in vitro* studies discovered cellular morphological changes that are consistent with acute, cytotoxic stress. Therefore, the observed tumor growth inhibition in the absence of CD8^+^ T cells expansion is likely due to a direct and transitory cytotoxic effect of berzosertib. Chemoproteomic selectivity targeting analysis has revealed that, while ATR remains the most potent target of berzosertib (> 10-fold), it engages multiple off-target proteins (35). In contrast, ceralasertib and elimusertib each displayed only single off-target engagement, with approximately 80-fold higher selectivity for ATR. This broader kinase engagement associated with berzosertib may contribute to its enhanced cytotoxicity, whereas the more selective inhibitors, ceralasertib and elimusertib, likely exert antitumor effects predominantly through ATR-dependent immunostimulatory mechanisms.

Previous studies using a similar treatment regimen have shown total CT26 rejection following rechallenge in complete responders (19). In our study, rechallenge of complete responder mice in the ceralasertib and elimusertib combination cohorts appeared to result in delayed tumor growth and deferred tumor rejection, not outright rejection. While the exact mechanism for this phenomenon was not fully explored, as this was not a primary objective of our experiment, a speculative reason may be that the failed tumor rejection upon rechallenge was due to our prior depletion of CD8^+^ T cells. The observation of delayed tumor growth may be due to incomplete depletion of tumor-specific memory CD8^+^ T cells, and/or due to the activity of other memory cells (e.g., CD4^+^, B-cells) that contributed to the antitumor response.

In our current studies, ATRi-induced immunostimulatory effects were dose dependent for ceralasertib and elimusertib, with the greatest response observed at 75 mg/kg and 40 mg/kg, respectively, while especially for elimusertib, an appreciable effect size is suggested even down to 1 mg/kg. Based on previous studies, these doses are associated with total exposures of 2,495 and 307 µg/mL•min, which are equivalent to unbound AUCs of 823 and 55.3 µg/mL•min, when accounting for the fraction of unbound drug in plasma (f^u,p^) in mice (20, 21) (**Suppl.Table 5**). Using this murine unbound AUC as a reference and applying the human f^u,p^ value of 22% and 4% for ceralasertib (**Suppl.Table 6**) and elimusertib, we were able to calculate a total human exposure needed of 3,741 and 1,383 µg/mL•min, respectively (36). Finally, using available PK parameters from clinical studies, we predicted clinical doses of 150 mg for ceralasertib and 270 mg for elimusertib, respectively, which would be required to elicit immune responses if combined with IR in humans (36, 37). Similarly, as berzosertib was associated with immune suppression, we predicted that a human equivalent dose of 115 mg/m^2^ would be needed for similar results in humans. Currently, ATRi are clinically administered in an intermittent, or pulsed, fashion to balance antitumor and toxic effects. Clinical trials of ceralasertib commonly incorporate a dosing regimen of 160-240 mg QD for 7-14 consecutive days of a 28 day cycle (38–41). The maximum tolerated dose (MTD) for elimusertib was determined to be 40 mg BID 3 days on/4 days off in a 21-day cycle (42). And the recommended phase 2 dose (RP2D) for berzosertib was 240 mg/m^2^ once- or twice-weekly in a 21-day cycle (43). Based on our predictions, we believe that ceralasertib clinical dosing is sufficient to enhance IR-mediated immune effects, under the assumption dosing is shortened to perhaps a few days only. Elimusertib clinical dosing is nearly 6-fold lower than the dose we predict to be associated with enhance IR-mediated immune effects, based on the highest mouse dose.

To summarize, our present work identifies that transient ATRi administration significantly enhances IR-induced antitumor activity. We report here, for the first time, that ATRi synergize with IR in a dose-dependent manner to increase (ceralasertib and elimusertib) or suppress (berzosertib) tumor antigen-specific CD8^+^ T cells in the DLN (**Figure 1**). When combined with IR, ATRi-enhanced immune stimulation was most pronounced with ceralasertib, however this effect was abolished with BID dosing and mimicked tumor antigen-specific CD8^+^ T cell populations previously observed with QD dosing of berzosertib. Given the differences in half-life between ATRi (ceralasertib: 1.75 h; elimusertib: 1.80 h; berzosertib: 6.12 h) this suggested that brief, ATR inhibition is required to be transient for antigen-specific CD8^+^ T cell expansion in the DLN (**Figure 2**). When combined with IR, two cycles of any ATRi treatment (ceralasertib, elimusertib, or berzosertib), significantly reduced tumor burden compared to IR alone (**Figure 3**). Although significant increases in survival and antitumor memory were only observed in mice receiving ceralasertib or elimusertib in combination with IR (**Figure 4)**, aligning with our immunostimulatory findings. The distinct immunomodulatory and cytotoxic effects observed *in vitro* with each ATRi (**Figure 5**) suggest additional mechanisms of actions for ceralasertib/elimusertib relative to berzosertib, potentially contributing to immune-mediate tumor shrinkage in addition to direct tumor cell killing, respectively, and offer an explanation as to why at least some antitumor benefit was observed with all 3 ATRi.

As targeted therapeutics continue to replace broad cytotoxics in anticancer regimens, rational combinations that exploit both DNA damage and immune system priming may offer promising approaches to improve patient outcomes. Our findings indicate that enhancement of IR antitumor activity is not a uniform class effect for ATRi and differences in ATRi half-life, persistence, and duration of exposure are critical factors in shaping therapeutic outcomes.

## 4 METHODS

### 4.1 Sex as a biological variable

While rodents can exhibit sex specific differences in absorption and elimination, the magnitude of these differences does not appear to translate to non-rodent species. Within this manuscript, we therefore performed the studies in the same sex (female) as where we originally performed our PK studies. Duplicating these efforts in male rodents would not contribute to a meaningful ultimate translation to humans and would merely result in an undesired duplication of non-vertebrate animal use. Mixing male and female rodents may result in increased variability in drug exposure within groups, with concomitant increased variability in outcomes, and decrease in power.

### 4.2 Cell lines and reagents

CT26 (CRL-2638) cells were purchased from ATCC (Manassas, VA) and cultured in RPMI-1640 medium with L-glutamine (BioWhittaker Inc., Walkersville, MD), containing 10% heat-inactivated FBS and 100 units of penicillin/mL and 100 µg/mL of streptomycin (Biofluids, BioSource, Rockville, MD) in an incubator with 5% C0^2^ and 95% humidity at 37 °C. Cells were checked for mycoplasma by IDEXX BioAnalytics (Westbrook, ME). Ceralasertib, elimusertib, and berzosertib were purchased from AdooQ Bioscience (Irvine, CA) and dosed as previously described (20–22). AH1 peptide (SPSYVYHQF) was diluted to a final concentration of 10 µg/mL in T Cell Media (RPMI, 10% FBS, 100 U/mL penicillin, and 100 mg/mL streptomycin).

### 4.3 Human plasma protein binding and bioanalysis

Ceralasertib protein binding in human plasma was performed *in vitro* using rapid equilibrium dialysis (RED) devices (Thermo Fisher Scientific, Waltham, MA). Freshly spiked human plasma (200 µL) and blank PBS (350 µL) were added to the sample and buffer chambers, respectively. Samples were incubated in triplicate at 37 °C for 24 h at 100, 1,000, and 10,000 ng/mL. Final plasma samples contained < 0.1% organic solvent.

Ceralasertib quantitation in human plasma was performed using an FDA-validated method as previously described (21). Briefly, an LC-MS/MS assay was implemented on a system consisting of an Agilent (Palo Alto, CA) 1290 Infinity II Autosampler and Binary Pump and a SCIEX (Concord, ON, Canada) 6500+ mass spectrometer. This included the use of [^2^H^4^]-ceralasertib as an isotopic internal standard. MRM channels were 413.2>334.0 for ceralasertib and 417.2.338.2 for [^2^H^4^]-ceralasertib. Chromatographic separation was achieved using a Phenomenex (Torrance, CA) Synergi Polar-RP 80A column (4 µm, 2mm x 50 mm) at ambient temperature with a gradient mobile phase consisting of mobile phase solvent A (water with 0.1% formic acid) and mobile phase solvent B (methanol with 0.1% formic acid). Retention times were 3.2 min for both ceralasertib and internal standard.

### 4.4 Mice and treatments

Specific pathogen-free female BALB/c mice (4-6 weeks of age) were purchased from Charles River (Wilmington, MA). Mice were allowed to acclimate to the University of Pittsburgh Animal Facility for at least 1 week before studies were initiated. To minimize exogenous infection, mice were maintained in microisolator cages and handled in accordance with the Guide for the Care and Use of Laboratory Animals (National Research Council, 2011) and on a protocol approved by the University of Pittsburgh IACUC. Ventilation and airflow in the animal facility were set to 12 changes/h. Room temperature was regulated at 72 ± 4 °F and the rooms were kept on automatic 12-h light/dark cycles. The mice received Prolab ISOPRO RMH 3000, Irradiated Lab Diet (PMI Nutrition International, Brentwood, MO) and water *ad libitium*. Throughout all studies, mice were routinely weighed and monitored for changes in health.

#### 4.4.1 Tumor

Mice were injected subcutaneously on the right hind flank with 1x10^6^ CT26 cells (ATCC, Manassas, VA). Cells were cultured in RPMI-1640 medium with L-glutamine (BioWhittaker Inc., Walkersville, MD), containing 10% heat-inactivated fetal bovine serum and 100 units of penicillin/mL and 100 µg/mL of streptomycin (Biofluids, BioSource, Rockville, MD) in an incubator with 95% air, 5% CO2, and 95% humidity at 37 °C. Cells were verified and checked for mycoplasma by IDEXX BioAnalytics (Westbrook, ME). Tumor growth was monitored twice weekly with a digital caliper, and tumor volume was calculated as volume = (length • width^2^)/2. Treatment was initiated once tumors reached approximately 90–150 mm. The survival endpoint was designated as tumor volume > 2,000 mm^3^.

For the IR +/- ATRi dose-response study, immobilized mice were exposed to 2 fractions of 2 Gy IR (6 mV photon energy, 2 cm field, 100 cm SSD) on days 1-2. Doses, route of administration, and schedule of ATRi for these studies were as follows: 75, 20, 7.5 or 2 mg/kg PO for ceralasertib; 40, 10, 4 or 1 mg/kg PO for elimusertib; and 2, 6, 20, or 60 mg/kg IV for berzosertib administered on days 1-3. For mice receiving combination treatments, IR was delivered approximately 30 min after ATRi administration. For each of these studies, mice were stratified into four treatment groups of: ATRi (ceralasertib, elimusertib, or berzosertib) alone, IR (2 gy x 2) alone, ATRi and IR combined, or respective ATRi vehicle control. The vehicle for ceralasertib was 10% DMSO, 40% propylene glycol, and 50% dH2O; the vehicle for elimusertib was 10% ethanol, 10% cremophor, and 80% saline; and the vehicle for berzosertib was 12% captisol adjusted to a pH of 4 with acetic acid. Each treatment group per study consisted of 5 mice. Each study was repeated to obtain a sample size of 10 per treatment group. Studies for each ATRi were conducted separately with independent IR alone and vehicle control groups.

Mice receiving radiation in the ceralasertib QD vs BID dosing study and the efficacy/survival study, were exposed to 2 fractions of 2 Gy IR on days 1-2 using an image-guided Precision SmART+ (225 kV, 20 mA energy) with a 0.3 mm Cu treatment filter and 10 mm circular collimator (Precision X-Ray, Madison, CT). Radiation was delivered at a 30-degree beam angle to mice in the prone position to minimize exposure of normal tissue to the beam field. For this study, mice also received ceralasertib 75 mg/kg PO QD or BID on days 1-3. Mice receiving ceralasertib BID were exposed to IR approximately 30 min after the initial dose followed by an 8-hour waiting period before the second dose.

#### 4.4.2 Tissue processing, staining, and flow cytometry

On day 9, CT26 tumors (dose-response study only) and DLNs were harvested from mice and placed into R10 media (RPMI/10% FBS). DLNs were mechanically dissociated between two frosted class slides and filtered through 70 μm cell strainers (Gibco, Thermo Fisher Scientific, Waltham, MA) to generate single-cell suspensions.

Tumor tissue was first injected with 1.5–2 ml digestion solution consisting of 50 μg/ml Liberase DL research grade (Roche) and 10 U/ml DNase I (Sigma-Aldrich, St. Loius, MO) in RPMI. After incubating 3–5 minutes at RT, tumors were finely chopped with a sterile razor blade and incubated in a total volume of 5 ml digestion solution for 15-20 minutes in an incubator set at 37 °C. Tumor pieces were then dissociated between frosted glass slides and filtered through 70 μm cell strainers as previously described. To generate single-cell suspensions, the solution was vortexed on a low speed for approximately 90 seconds and filtered again through a new 70 μm cell strainer. RBCs were lysed with 1 mL RBC lysis buffer (150 mM NH^4^Cl, 10 mM NaHCO^3^, 0.1 mM EDTA at pH 8.0) for approximately 10 seconds at RT followed by quenching with 4 mL RPMI. Cell suspensions were counted using a Scepter handheld counter (MilliporeSigma, Burlington, MA) and were seeded at 1.5 × 10^6^ cells in 96-well round-bottom plates for staining. Plated cells were first subjected to Fc receptor block with 0.5 µg anti-CD16/32 antibody (TruStain FcX™, BioLegend, San Diego, CA) in FSC buffer (2% FBS/1x PBS) for 10 min at 4°C. CT26 tumor antigen-specific CD8^+^ T cells were then labeled with 10 µL PE-labeled H-2Ld–SPSYVYHQF (AH1 peptide) Pro5 MHC Class I Pentamer (ProImmune, Oxford, UK) in FSC buffer (60 µL total volume) for 10 min at RT. With each experiment, PE-labeled HLA-A*2:01 Negative Control Pro5 MHC Class I Pentamer (ProImmune, Oxford, UK) and lymph nodes from naïve, no tumor, negative control mice were included in appropriate control wells. An antibody cocktail against surface antigens was prepared in FCS buffer, added to cells, and incubated for 15 min at 4°C. The cocktail included the following antibodies: anti-CD8 (Mouse) mAb-Alexa Fluor® 647 (MBL International, Woburn, MA), γδ T-Cell Receptor Hamster anti-Mouse, Brilliant Violet 786 (Clone: GL3) BD Optibuild™ (BD Biosciences, San Jose, CA), Alexa Fluor® 488 anti-mouse CD45, Brilliant Violet 785™ anti-mouse CD4, Alexa Fluor® 488 anti-mouse CD19, and Brilliant Violet 785™ anti-mouse CD335 (NKp46) (all from Biolegend, San Diego, CA). After washing with FCS buffer, cells were stained with eFluor™ 780 viability dye in 1x PBS and allowed to incubate 10 min at 4 °C. Cells were then fixed for 1 h at RT in 1x FluorFix (eBioscience, Thermo Fisher Scientific, Waltham, MA), washed twice with FCS buffer, and stored in FSC buffer at 4 °C in the dark overnight prior to acquisition.

Uncompensated data was collected the following morning using an LSRFortessa cytometer with FACSDiva software (both BD Biosciences, San Jose, CA). Compensation and subsequent data analyses were performed with FlowJo v10 software (FloJo LLC, Ashland, OR). Single stained DLN samples with matching unstained cells or single stained OneComp eBeads (Thermo Fisher Scientific, Waltham, MA) were used for single-color compensation controls. Fluorescence-minus-one (FMO) controls were utilized when appropriate to empirically set gating. For each experiment, gates were applied uniformly for all samples, including determination of the pentamer gate by FMO controls which were stained with the entire antibody staining panel, except pentamer.

#### 4.4.3 Measurement of cytokines/chemokines and morphological analysis of CT26 cells

CT26 cells were maintained in complete DMEM supplemented with 10% FBS and penicillin/streptomycin, and cultured at 37 °C with 5% CO^2^. Experiments were conducted within 11 generations and cells were checked periodically for mycoplasma contamination by PCR.

For measurement of IFN-β, CXCL10, and CCL2, CT26 cells were seeded in a 60 mm dish (70,000 cells/dish) overnight followed by treatment with one of the following ATRi at IC^25^ concentrations: ceralasertib (5 µM), elimusertib (0.25 µM), or berzosertib (0.5 µM). After 24 h, cells were collected using 300 µL of TRIzol (Invitrogen, Thermo Fisher Scientific, Waltham, MA) and processed for RNA isolation using the using the Direct-zol RNA Miniprep Kit (Zymo Research, Irvine, CA) according to the manufacturer’s instructions. Total RNA, (100 ng) was reverse transcribed using Lunascript RT master mix (New England Biolabs, Ipswich, MA) and diluted 10-fold. Each PCR reaction was run as technical duplicates using 5 µL of Luna Universal qPCR master mix (New England Biolabs, Ipswich, MA), 1 µL of each 2.5 µM primer, and 3 µL of diluted cDNA. The PCR program was run according to the manufacturer’s suggestions and quantification was performed using Quantstudio software (Thermo Fisher Scientific, Waltham, MA). The forward and reverse primer sequences were as follows: *GAPDH* forward primer: 5′-CTCTGGAAAGCTGTGGCGTGATG-3′ and *GAPDH* reverse primer: 5 `′` - ATGCCAGTGAGCTTCCCGTTCAG-3 `′` . *IFN-β* forward primer: 5 `′` - AAGAGTTACACTGCCTTTGCCATC-3 `′` and *IFN-β* reverse primer 5 `′` - CACTGTCTGCTGGTGGAGTTCATC-3`′` . *CXCL10* forward primer: 5’-ACTGCATCCATATCGATGAC and *CXCL10* reverse primer: 5′-TTCATCGTGGCAATGATCTC-3′. *CCL2* forward primer: 5’-GCAGCAGGTGTCCCAAAGA-3′ and *CCL2* reverse primer: 5’-TTGGTTCCGATCCAGGTTTTT-3′.

For imaging of CT26 morphology following *in vitro* treatment with ATRi, cells were seeded into a black, flat-bottom 96-well plate (2,500 cells/well). After 16 h, cells were treated with one of the following ATRi at increasing concentrations: ceralasertib (0.15, 0.5, 1.5, 5, or 50 µM), elimusertib (0.015, 0.05, 0.15, 0.5, or 5 µM), berzosertib (0.15, 0.5, 1.5, 5, or 50 µM). After 24 h, cells were washed 1x with PBS and fixed with 2% paraformaldehyde (PFA) in the dark for 15 min. Cells were washed 3x with PBS, cell nuclei was stained with NucBlue ReadyProbes reagent (Thermo Fisher Scientific, Waltham, MA) in the dark for 1 h. Cell images were captured in brightfield and fluorescence channels using 4x PL FL Phase objective on a Nikon Fluorescence Microscope.

### 4.5 Statistics

Tumor antigen-specific CD8+ T cell response to each ATRi drug in combination with a fixed IR regimen was analyzed across treatment groups using a linear model with ATRi dose as a continuous variable and the IR alone group receiving 0 mg/kg of ATRi. Only groups receiving IR were included in the model. Group differences in the percentage of CD45+CD8+ T cells in the DLN were assessed using a Kruskal-Wallis test with Dunn’s post-hoc correction for multiple comparisons. BID vs QD dosing of ceralasertib or elimusertib plus IR was evaluated by a generalized linear model. The percentage change in tumor volume on day 26 relative to day 1 was assessed by one-way ANOVA with Dunnett’s post-hoc correction for multiple comparisons. Tumor growth over time was analyzed by a linear mixed-model to adjust for repeated measures over time. Tumor volume was log-transformed to achieve linearity and a random intercept was specified for each mouse along with fixed effects for treatment group and time. Due to the complete response for several mice in the ceralasertib and elimusertib groups, these mice were excluded from the tumor growth model due to inherently different tumor growth curves for mice whose tumor volume achieved and remained at 0. These responses are noted in the corresponding figures and growth curves are included for completeness. Growth factor was defined per treatment group (excluding complete responders) and represents the base of the exponential function for change in volume per day. Survival curves were estimated by the Kaplan-Meier method with mice surviving until study termination censored at day 58. One additional mouse was censored at day 16 due to a dosing issue unrelated to disease or treatment. Number at risk was defined as total number of non-censored surviving mice per treatment group at each specified time. Survival across all groups was compared by a log-rank test. A Cox proportional hazard model was fit for all IR-treated groups and pairwise comparisons were made to the “IR alone” control group. In general, analysis of experimental repeats found no significant batch effects, therefore data were combined across repeats for further analysis without controlling for batch. Prior to initiation of the IR ± ATRi efficacy and survival studies, group differences in total body weight and starting tumor volume (post-stratification and randomization) were assessed using a Kruskal-Wallis test. Analysis of GAPDH-normalized gene expression changes were assessed by Friedman test with Dunn’s post-hoc correction for multiple comparisons. Confidence intervals are reported at the 95% significance level and p-values below 0.05 were considered statistically significant for all tests unless otherwise specified. Brackets are shown only for comparisons that were statistically significant unless otherwise specified. Statistical analysis was performed using R version 4.4.1 or GraphPad Prism (version 10.2.2).

### 4.6 Study approval

All experimental procedures were approved by the University of Pittsburgh Animal Care and Use Committee and performed in accordance with relevant guidelines and regulations.

## AUTHOR CONTRIBUTIONS

JJD: Writing – review & editing, Writing – original draft, Visualization, Methodology, Investigation, Formal analysis, Data curation, Conceptualization. BFK: Writing – review & editing, Writing – original draft, Visualization, Methodology, Investigation, Formal analysis, Data curation, Conceptualization. FPV: Writing – review & editing, Writing – original draft, Visualization, Methodology, Investigation, Formal analysis, Data curation, Conceptualization. PP: Writing – review & editing, Visualization, Methodology, Investigation. JG: Writing – review & editing, Methodology, Data curation. KLC: Writing – review & editing, Formal analysis, Data curation, Conceptualization. MT: Writing – review & editing, Methodology, Investigation. MdM: Writing – review & editing, Methodology, Investigation. NMI: Writing – review & editing, Methodology, Investigation. MJB: Writing – review & editing, Methodology, Data curation. DAC: Writing – review & editing, Resources, Methodology, Investigation, Conceptualization. CJB: Writing – review & editing, Resources, Methodology, Investigation, Funding acquisition, Formal analysis, Data curation, Conceptualization. JHB. Writing – review & editing, Writing – original draft, Supervision, Resources, Visualization, Methodology, Investigation, Funding acquisition, Formal analysis, Data curation, Conceptualization.

The authorship of co–second authors was determined to reflect their substantial contribution to the study, while acknowledging varying degrees of involvement in the collaborative work. The order was decided according to the first date of participation in the project.

## ACKNOWLEDGEMENTS

The authors would like to thank Dr. Ron Lalonde, chief physicist, UPMC Shadyside Hospital, for providing access to and supervising the use of the clinical irradiator. This project was also funded, in part, under a CURE Grant with the Pennsylvania Department of Health. The Department specifically disclaims responsibility for any analyses, interpretations, or conclusions. JJD was also supported, in-part, by a scholarship from the Community Foundation of Warren County (PA). BFK was also supported, in-part, by a fellowship from the American Foundation for Pharmaceutical Education. This project used the UPMC Hillman Cancer Center and shared resources supported, in-part, by award P30CA047904.

## Abbreviations

DDR: DNA damage response
ATR: Ataxia telangiectasia and Rad3-related
ATRi: Ataxia telangiectasia and Rad3-related inhibitor
IR: Ionizing radiation
CTL: Cytotoxic T lymphocyte
DLN: Draining lymph node
PK: Pharmacokinetic
QD: Once daily
TIL: Tumor infiltrating lymphocyte
BID: Twice daily
αCD8: Anti-CD8 antibody
TME: Tumor microenvironment
NK: Natural killer
ISG: Interferon-stimulated gene
^Fu,p^: Fraction unbound in plasma
MTD: Maximum tolerated dose
RP2D: Recommended phase 2 dose

