## SUPPLEMENTAL for "Transient ATR inhibition following ionizing radiation enhances immune-mediated antitumor response and survival"

*Authorship note: BFK and FPV are co-second authors

Correspondence:

Jan H. Beumer, PhD, PharmD, DABT

Department of Oncology

Johns Hopkins University School of Medicine and Johns Hopkins Sidney Kimmel Cancer Center

1650 Orleans St, Room 1M52, Baltimore, MD 21287, USA

Conflict-of-interest statement: JHB reports spouse salary from AstraZeneca.

### SUPPLEMENTAL TABLES

**Suppl.Table 1. Pairwise comparisons of antigen-specific $CD8^+$ T cell counts between QD or BID ATRi and IR alone cohorts.**

| Comparison | Ceralasertib 75 mg/kg | Elimusertib 40 mg/kg |
|---|---|---|
| IR+QD vs IR Alone | p=0.081 | p=0.127 |
| IR+BID vs IR Alone | p=0.144 | p=0.691 |
| IR+QD vs IR+BID | p=0.742 | p=0.061 |

P-values obtained from a generalized linear model fit with treatment group as a categorical variable. Values unadjusted for multiple comparisons.

**Suppl.Table 2. Pairwise comparisons of antigen-specific CD8$^+$ T cell counts between QD or BID ceralasertib (75 mg/kg) and IR alone cohorts following targeted IR.**

| Comparison | p-value |
|---|---|
| IR+QD vs IR Alone | < 0.001 |
| IR+BID vs IR Alone | 0.963 |
| IR+QD vs IR+BID | < 0.001 |

P-values obtained from a generalized linear model fit with treatment group as a categorical variable. Values unadjusted for multiple comparisons.

**Suppl.Table 3. Number of mice at risk per group during the IR ± ATRi survival study**

| Treatment Group | Day | | | | | |
|---|---|---|---|---|---|---|
| | 10 | 20 | 30 | 40 | 50 | 60 |
| No Drug Control | 9 | 4 | 0 | 0 | 0 | 0 |
| IR Alone | 9 | 7 | 4 | 0 | 0 | 0 |
| IR + 75 mg/kg ceralasertib | 10 | 8 | 8 | 5 | 4 | 0 |
| IR + 10 mg/kg elimusertib | 10 | 9 | 5 | 4 | 3 | 0 |
| IR + 20 mg/kg berzosertib | 10 | 9 | 4 | 1 | 0 | 0 |

**Suppl.Table 4. Results from the Cox proportional hazards model fit for all IR treated groups.**

| Comparison | Hazard Ratio vs IR Alone | 95% CI LL | 95% CI UL | P-value |
|---|---|---|---|---|
| IR+75 mg/kg Ceralasertib vs IR Alone | 0.22 | 0.08 | 0.65 | 0.006 |
| IR+10 mg/kg Elimusertib vs IR Alone | 0.33 | 0.12 | 0.93 | 0.037 |
| IR+40 mg/kg Berzosertib vs IR Alone | 0.72 | 0.29 | 1.78 | 0.472 |

All comparisons made to IR alone control. LL: lower limit. UL: upper limit.

**Suppl.Table 5. Murine total and unbound ATRi exposure scaled up to predict dose required to achieve optimal human exposure.**

| | Murine | | | | Human | | |
|---|---|---|---|---|---|---|---|
| | Dose Required for Immune Effect[a] | Total $AUC_{inf}$ | Fraction Unbound in Plasma | Unbound AUC[b] | Fraction Unbound in Plasma | AUC Needed to Achieve Murine Unbound AUC | Dose Required to Achieve Optimal AUC |
| ATRi | mg/kg | µg/ml*min | % | µg/ml*min | % | µg/ml*min | mg / $mg/m^2$ |
| Ceralasertib | 75 | 2,495 | 33 | 823 | 22 | 3,741 | 150 mg |
| Elimusertib | 40 | 307 | 18 | 55.3 | 4 | 1,383 | 270 mg |
| Berzosertib | 60 | 2,124 | 3 | 63.7 | 3 | 2,123 | 115 $mg/m^2$ |

[a]stimulatory or suppressive (when combined with IR)

[b]assuming equilibrium

**Suppl.Table 6. Ceralasertib protein binding in human plasma.**

| Concentration (ng/mL) | $F_{u,p}$ (%) | CV (%) |
|---|---|---|
| 100 | 0.244 | 3.77 |
| 1,000 | 0.198 | 9.41 |
| 10,000 | 0.231 | 11.4 |

N=3/concentration. $F_{u,p}$: fraction unbound in plasma

### SUPPLEMENTAL FIGURES

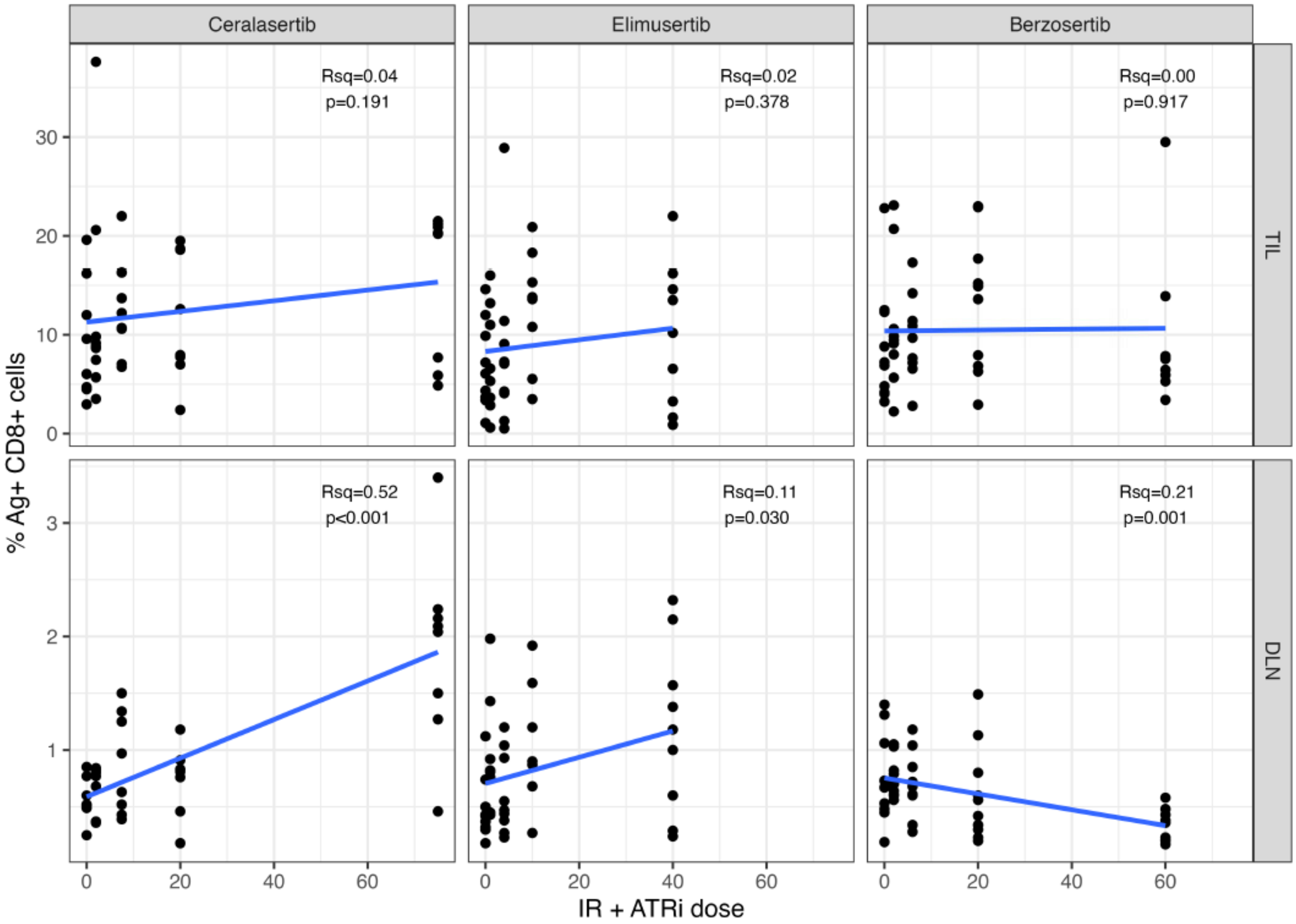


**Suppl.Fig. 1. Dose-response relationships between dose (IR + ATRi) and percentage of antigen-specific CD8+ T cells in tumor (TIL) and the DLN across ATRi treatment cohorts. Data represented as individual data from each treatment group receiving IR + ATRi. Significance reported if *$p < 0.05$* by linear model. Linear regression lines are shown in blue. Rsq: R-squared (coefficient of determination) value from linear model.**

DLN

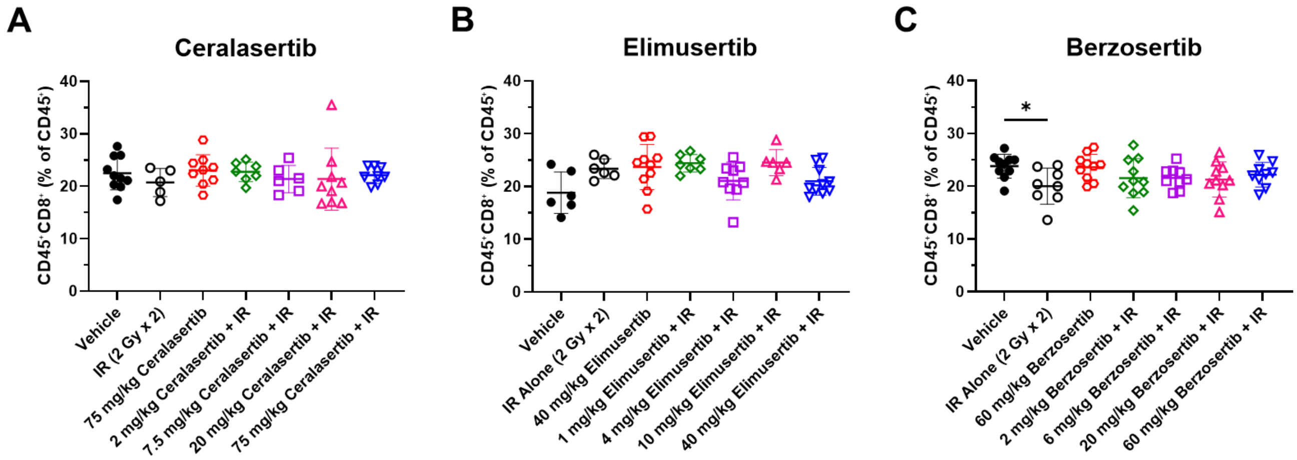


**Suppl.Fig. 2. The percentage of CD45$^+$CD8$^+$ T cells in mice receiving vehicle controls, IR alone, or A) ceralasertib B) elimusertib C) berzosertib alone or at escalating doses in combination with IR. Data represented as mean (bold line) and standard deviation. *P < 0.05, by Kruskal-Wallis with Dunn's post-hoc multiple comparisons test.**

DLN

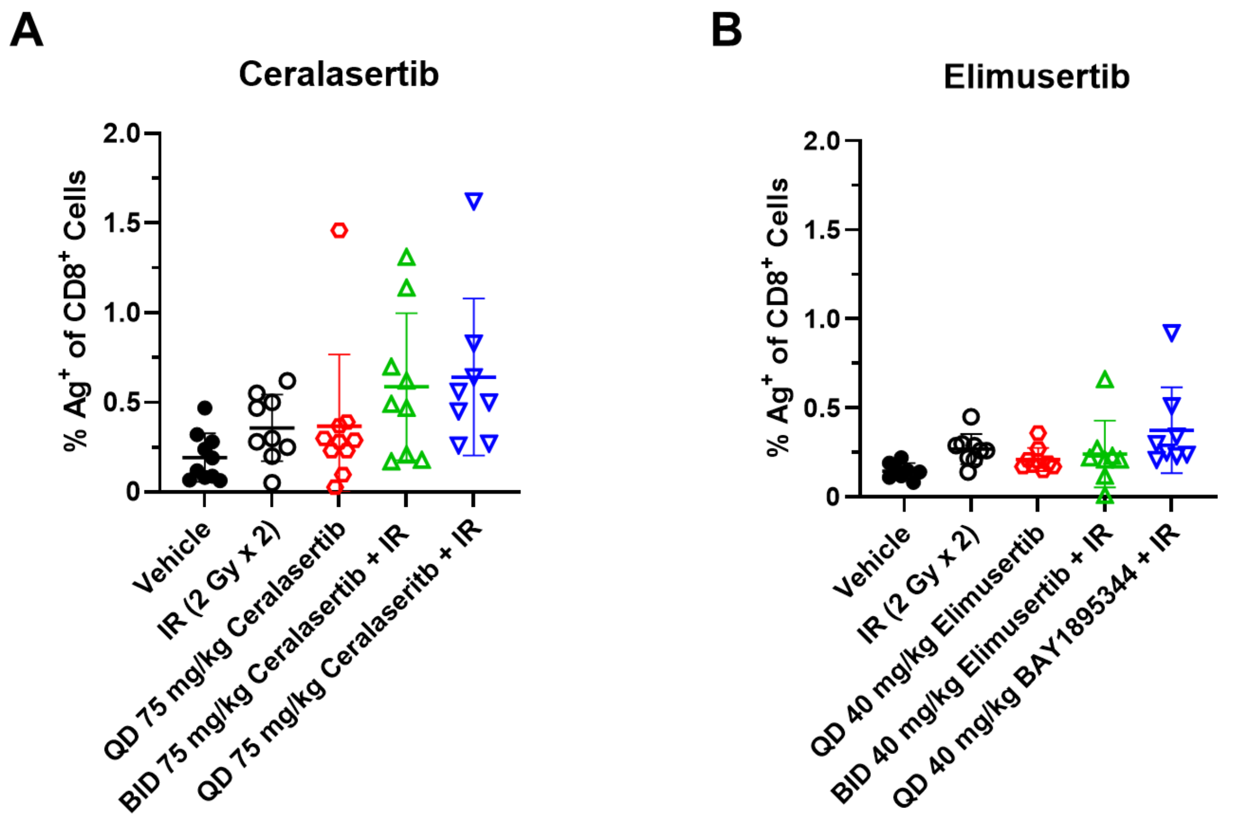


**Suppl.Fig. 3. The percentage of CT26 antigen-specific CD8$^+$ T cells following treatment with A) ceralasertib or B) elimusertib during the QD vs BID ATRi in combination with IR treatment studies. Data represented as mean (bold line) and standard deviation. No statistically significant differences were observed between treatment groups, as determined by fitting data to a generalized linear model.**

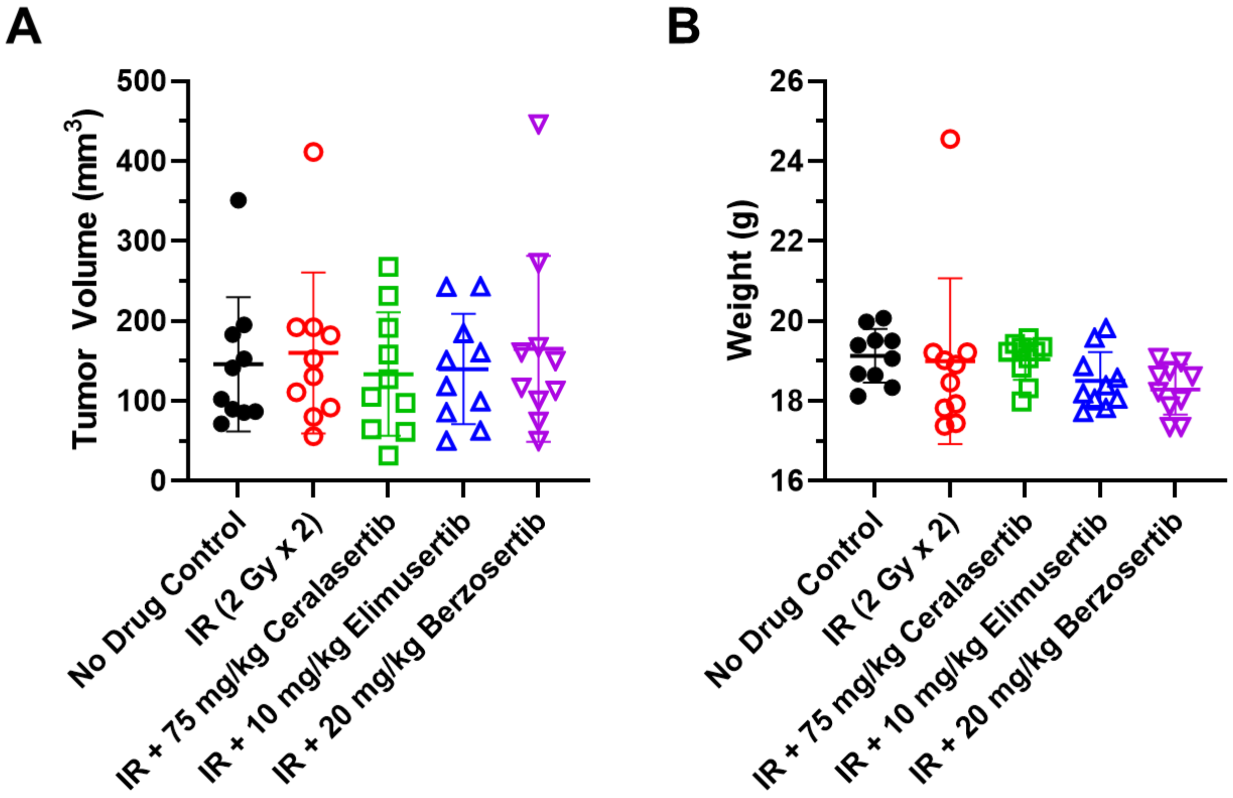


**Suppl.Fig. 4. Stratification of A) total body weight and B) baseline tumor volume prior to initialization of the efficacy and survival studies of ATRi in combination with IR. Data represented as mean (bold line) and standard deviation. No statistically significant differences were observed between groups, as determined by a nonparametric Kruskal-Wallis test.**

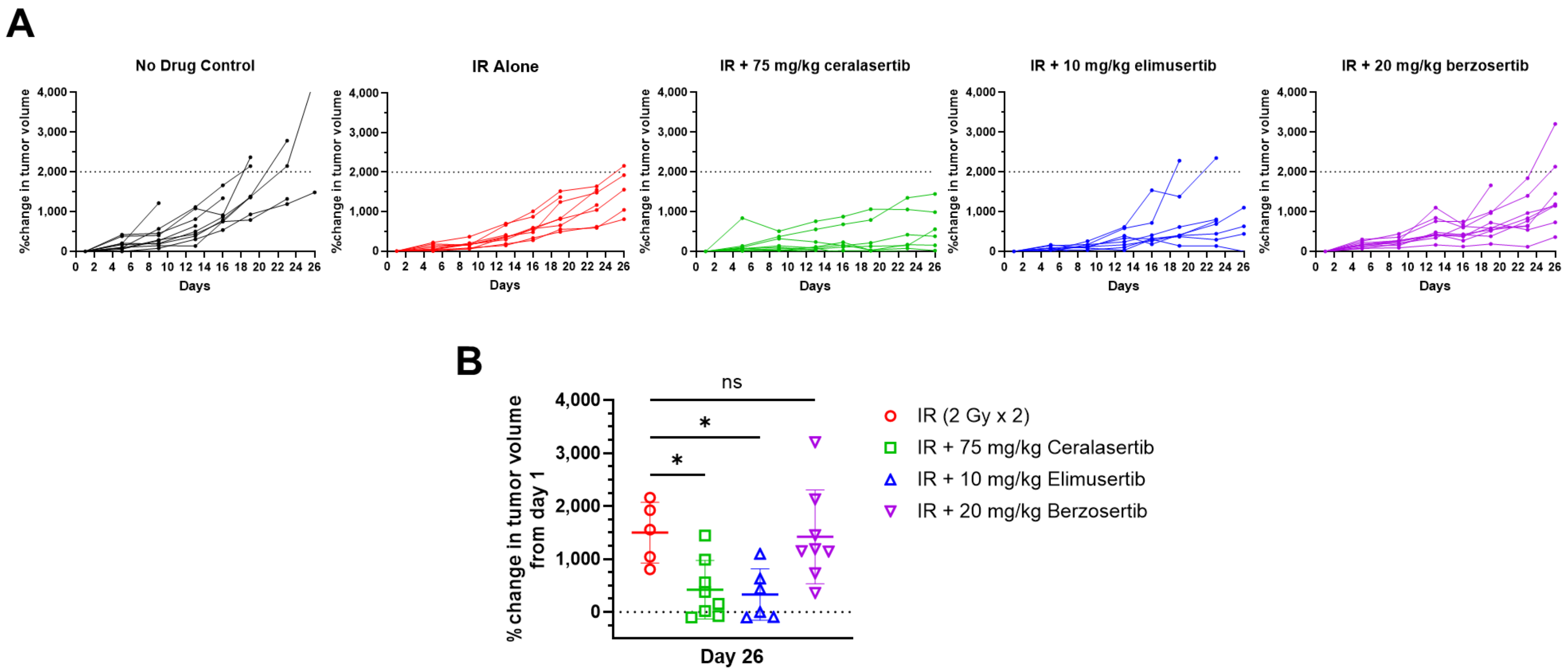


**Suppl.Fig. 5. The A) total percent change in tumor volume for all treatment cohorts at day 26 and B) percent change in tumor volume at day 26 for all treatment cohorts in individual mice relative to day 1. Data represented as mean (bold lines) and standard deviation. *$P < 0.05$ by one-way ANOVA with Dunnett's multiple-comparison test with the IR alone (2 Gy x 2) group serving as the control. NS: not statistically significant.**

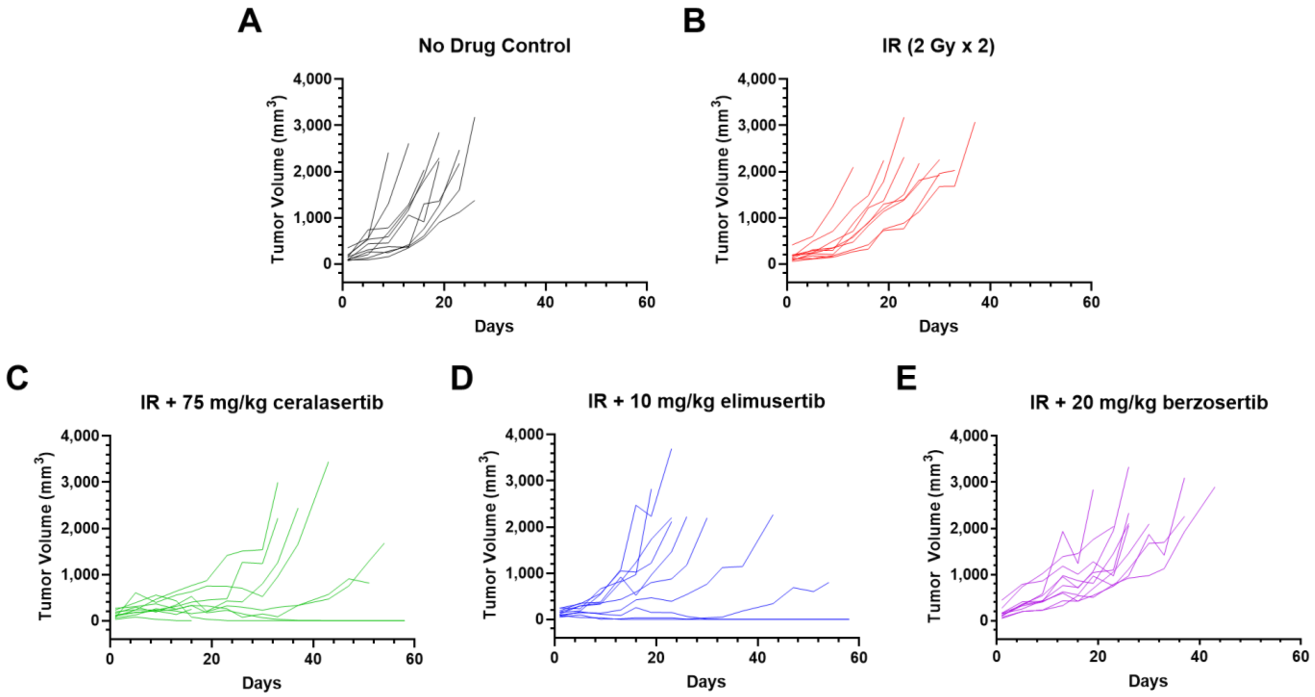


**Suppl.Fig. 6. Absolute tumor volumes for A) No Drug Control, B) IR (2 Gy x 2), C) 75 mg/kg ceralasertib + IR, D) 10 mg/kg elimusertib + IR, and E) 20 mg/kg berzosertib + IR cohorts during the course of the efficacy and survival studies of ATRi in combination with IR.**
